# Decoupling SARS-CoV-2 ORF6 localization and interferon antagonism

**DOI:** 10.1101/2021.12.06.471415

**Authors:** Hoi Tong Wong, Victoria Cheung, Daniel J. Salamango

## Abstract

Like many pathogenic viruses, SARS-CoV-2 must overcome interferon (IFN)-mediated host defenses for infection establishment. To achieve this, SARS-CoV-2 deploys overlapping mechanisms to antagonize IFN production and signaling. The strongest IFN antagonist is the accessory protein ORF6, which localizes to multiple membranous compartments, including the nuclear envelope, where it directly binds the nuclear pore components Nup98-Rae1 to inhibit nuclear translocation of activated STAT1/IRF3 transcription factors. However, a direct cause-and-effect relationship between ORF6 localization and IFN antagonism has yet to be explored experimentally. Here, we use extensive mutagenesis studies to define the structural determinants required for steady-state localization and demonstrate that mis-localized ORF6 variants can still potently inhibit nuclear trafficking and IFN signaling. Additionally, expression of a peptide that mimics the ORF6/Nup98 interaction domain robustly inhibited nuclear trafficking. Furthermore, pharmacologic and mutational approaches combined to suggest that ORF6 is likely a peripheral-membrane protein, opposed to being a transmembrane protein as previously speculated. Thus, ORF6 localization and IFN antagonism are independent activities, which raises the possibility that ORF6 may have additional functions within membrane networks to enhance virus replication.

## INTRODUCTION

Severe acute respiratory syndrome coronavirus-2 (SARS-CoV-2) is a novel coronavirus responsible for causing the coronavirus disease 2019 (COVID19) pandemic, which continues to be a persistent problem worldwide despite the availability of highly effective vaccines. A major barrier to wide-spread immunity has been the emergence of novel variants that exhibit significantly increased transmission rates. Pathogenicity is further enhanced through the activities of several viral proteins that suppress type I interferon (IFN) production and signaling. For instance, SARS-CoV-2 NSP1, NSP6, NSP13, ORF6, ORF7b, ORF8, and Nucleoprotein all antagonize IFN signaling to various degrees, with ORF6 exhibiting the strongest inhibition (Lei et al., 2020; Miorin et al., 2020; Xia et al., 2020).

ORF6 is a 7 kilodalton accessory protein required for optimal virus replication *in vitro* and *in vivo*—likely through its ability to potently suppress IFN signaling (Silvas et al., 2021; Zhao et al., 2009; Zhou et al., 2010). Several studies have demonstrated that ORF6 induces cytoplasmic accumulation of importinα and importinβ, which subsequently inhibits nuclear translocation of activated STAT1 and IRF3 transcription factors (Frieman et al., 2007; Lei et al., 2020; Miorin et al., 2020). Additional mechanistic clarification was recently provided through proteomics studies that identified the Nup98-Rae1 nuclear pore complex as high-confidence ORF6 interactors (Gordon et al., 2020; Miorin et al., 2020). Further investigation revealed that a direct interaction between the C-terminal tail of ORF6 and the C-terminal domain of Nup98-Rae1 impairs docking of cargo/receptor complexes to inhibit nuclear trafficking (Addetia et al., 2021; Kato et al., 2021; Miorin et al., 2020).

While suppression of antiviral innate immune signaling is believed to be its primary function, ORF6 also packages into nascent viral particles and induces membrane structures resembling viral replication compartments (Huang et al., 2007; Zhou et al., 2010). These activities coincide with ORF6 subcellular distribution, as it localizes to several membranous compartments during infection and when exogenously expressed in several different cell types (Gunalan et al., 2011; Kumar et al., 2007). Both native and epitope tagged ORF6 proteins colocalize with markers for the Golgi apparatus (Kato et al., 2021; Kopecky-Bromberg et al., 2007; Lei et al., 2020; Miorin et al., 2020; Xia et al., 2020; Zhou et al., 2010), endoplasmic reticulum (Kopecky-Bromberg et al., 2007; Lee et al., 2021; Lei et al., 2020; Zhou et al., 2010), endosomes (Gunalan et al., 2011; Kumar et al., 2007; Lee et al., 2021), and nuclear envelope (Addetia et al., 2021; Kato et al., 2021; Miorin et al., 2020), which is consistent with the current paradigm that ORF6 is a transmembrane protein capable of lateral diffusion within membranous networks (Netland et al., 2007; O’Keefe et al., 2021; Zhou et al., 2010). Taken together, these observations lend to an appealing model where ORF6 localization at the nuclear envelope facilitates a direct interaction with Nup98-Rae1 to subsequently inhibit nuclear trafficking; however, this direct cause-and-effect relationship has yet to be examined experimentally.

Here, we investigate the link between ORF6 localization and IFN antagonism. Through an extensive panel of truncation and single amino acid substitution mutants in combination with pharmacologic experiments, we demonstrate that ORF6 associates with membranous compartments through two distinct structural determinants and is most likely a peripheral-membrane protein. The first determinant resides within amino acid residues ^18^IMRTFKV^24^ and is required for Golgi retention. The second determinant encompasses two putative amphipathic helices required for maintaining steady-state localization and membrane association. Importantly, mis-localized ORF6 variants potently induced cytoplasmic accumulation of importinα, inhibited nuclear translocation of activated STAT1, and suppressed IFN signaling. In further support of these observations, a peptide inhibitor that contains the ORF6/Nup98 interaction motif robustly blocked nuclear accumulation of importinα. Taken together, these results demonstrate that membrane association of ORF6 is dispensable for interferon antagonism, which raises the possibility that ORF6 may have additional functions within membrane networks.

## RESULTS AND DISCUSSION

### Identification of ORF6 localization determinants

Native and epitope tagged SARS-CoV-2 ORF6 proteins localize to several membranous organelles; however, the determinants that dictate subcellular distribution have yet to be thoroughly investigated. To identify which protein region mediates steady-state localization, a panel of ORF6-mCherry truncation mutants were generated and co-expressed in HeLa cells with markers for the ER and Golgi apparatus (**Fig. 1**). A computational model was used to generate truncations that minimized perturbations to putative structural elements as a structure has yet to be solved (**Fig. 1A**). Because the C-terminal region of ORF6 is required for interacting with Nup98-Rae1 and for inducing cytoplasmic accumulation of importinα (Frieman et al., 2007; Miorin et al., 2020), we reasoned that any *cis-*acting localization determinants reside upstream of this protein/protein interaction domain. As expected, expression of amino acid residues 1-47 in HeLa cells resulted in localization comparable to wild-type, while expression of residues 48-61 had no discernable localization pattern (**Fig. 1B**). Unexpectedly, when we further examined the 1-47 segment for *cis*-acting residues that dictate localization, we were surprised to observe two distinct localization patterns. Expression of the first half of this segment (amino acid residues 1-23) resulted in partial colocalization with the Golgi marker, while expression of the second half (amino acid residues 24-47) resulted in localization to an organelle distinct from the ER (**Fig. 1B**, 1-23 and 24-47, merge). When the 1-23 segment was further truncated to residues 1-17, no localization pattern was observed, suggesting that Golgi retention is partially mediated by amino acid residues ^18^IMRTFKV^24^ (**Fig. 1B**).

**Figure 1.**
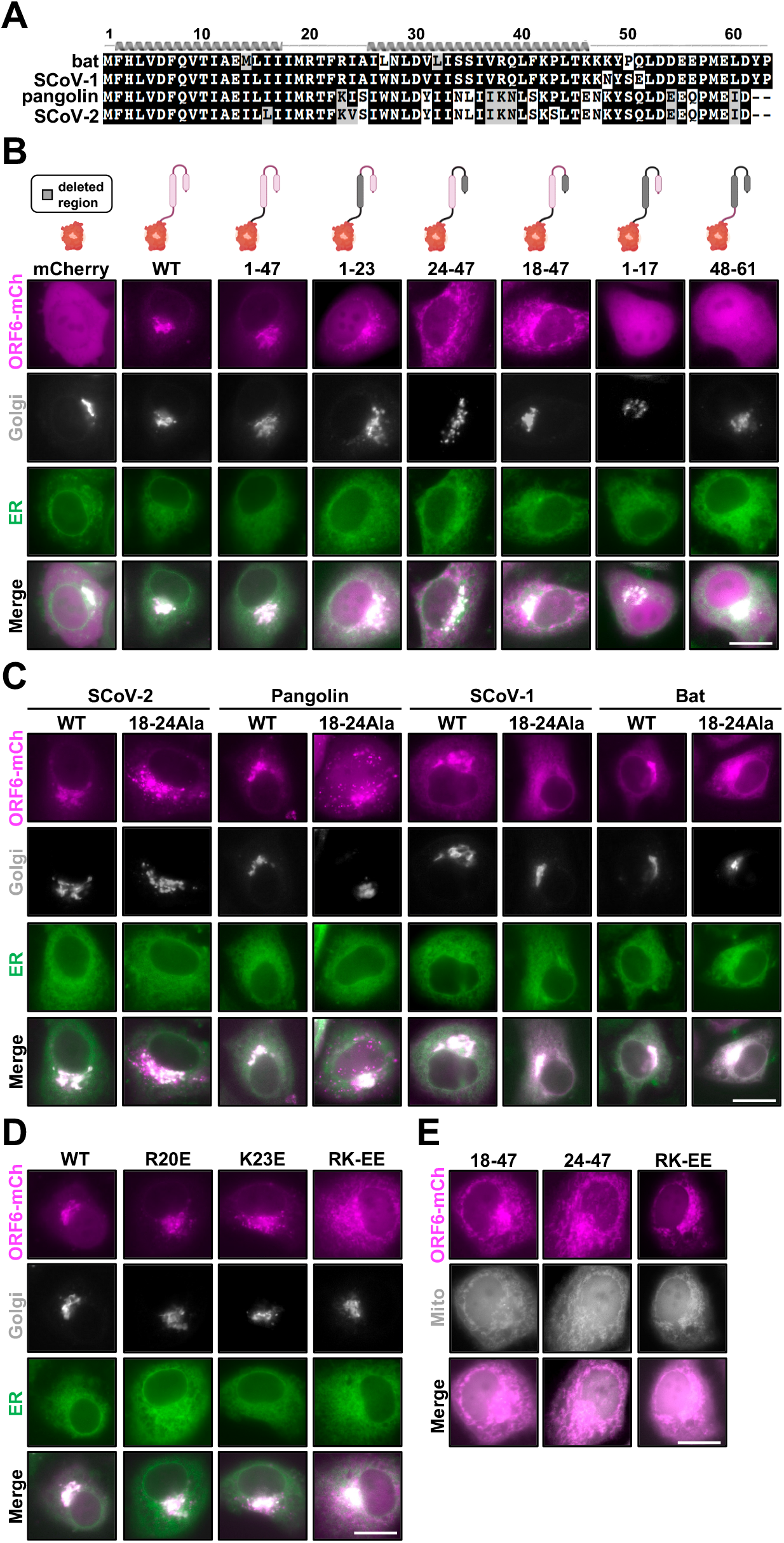
ORF6 truncation mutants reveal key localization determinants. (**A**) ORF6 amino acid alignment of the indicated species with secondary structure prediction. (**B**-**E**) Representative live cell fluorescence microscopy images of the indicated mCherry-tagged ORF6 proteins expressed with the indicated subcellular markers in HeLa cells.

Because the 1-23 fragment only partially recapitulated Golgi retention, we reasoned that the putative helix within residues 24-47 is required for maintaining steady-state localization, since when expressed alone it non-specifically associated with a membranous compartment. Therefore, we generated a truncation construct that expressed amino acid residues 18-47 anticipating it would mimic wild-type distribution; instead, this mutant localized to the ER-adjacent organelle, suggesting that the putative helix contained within the 1-17 fragment also contributes to steady-state localization (**Fig. 1B**, 1-47 and 18-47, merge). These observations suggested that ORF6 maintains steady-state localization through at least two distinct determinants, a longer protein component from residues 1-47 that mediates steady-state membrane associations, and a second region from ^18^IMRTFKV^24^ that dictates Golgi retention. To test this model, we initially focused on ^18^IMRTFKV^24^ to determine if it harbors a Golgi retention motif. Substitution of ^18^IMRTFKV^24^ to alanine in full length ORF6 not only disrupted Golgi accumulation but induced freely diffuse intracellular puncta (**Fig. 1C**, 18-24Ala). Further investigation into the conservation of this motif revealed that SARS-CoV-1, bat, and pangolin ORF6 proteins also require this motif to facilitate Golgi retention (**Fig. 1C**). Of note, SARS-CoV-1 and bat 18-24Ala mutants did not form intracellular puncta as compared to SARS-CoV-2 and pangolin ORF6, but exhibited strong colocalization with the ER marker, suggesting there is an inherent difference in the way these proteins associate with membranes (**Fig. 1C**, merge). It is possible this difference is attributable to the putative helix from residues 24-47, as there is only ∼50% amino acid identity within this region across species (**Fig. 1A**).

While no strict consensus motif has been defined for Golgi retention, numerous Golgi resident proteins maintain steady-state localization through motifs enriched with positively charged amino acid residues (Tu et al., 2012; Wang et al., 2020); and interestingly, ^18^IM<u>R</u>TF<u>K</u>V^24^ contains arginine and lysine residues at positions 20 and 23, respectively. To test the contribution of these residues to Golgi retention, we generated single and double amino acid substitution mutants at these positions and assessed localization patterns. Independently exchanging these residues for glutamate had little impact on ORF6 localization when expressed in HeLa cells; however, the RK20,23EE double mutant lost Golgi retention and localized to the ER-adjacent organelle, raising the possibility that electrostatic interactions may facilitate ORF6 targeting to the Golgi (**Fig. 1D**, merge). This speculation is intriguing given that the Golgi membrane is enriched with phosphatidylinositol-4-phosphate lipids, which unlike phosphatidylcholine, ethanolamine, and serine, contain negatively charged headgroups.

Because several of the ORF6 mutants strongly localized to an ER-adjacent organelle, we co-expressed relevant ORF6 mutants with markers for the most likely candidates to identify this compartment (*i*.*e*., mitochondria and ER-Golgi-intermediate compartment). Interestingly, all ER-adjacent mutants exhibited significant colocalization with the mitochondria, but not with the ER-Golgi-intermediate compartment (**Fig. 1E**, and data not shown).

### SARS-CoV-2 ORF6 is likely a peripheral-membrane protein

While fine-mapping ORF6 localization determinants, several of the mutants exhibited colocalization with the mitochondria (SARS-CoV-2 18-24, 24-47, and RK-EE) or ER (SARS-CoV-1 and bat 18-24Ala), raising the possibility that a second determinant drives association with membranous compartments (**Fig. 1**). Because it has been reported that transmembrane domains of Golgi resident proteins contribute to steady-state localization (Banfield, 2011; Hu et al., 2011; Wang et al., 2013), we reasoned that the putative ORF6 alpha helices identified in the computational model contribute to localization. To explore this possibility, we closely examined the amino acid composition of these helices looking for clues as to how they might associate with membranous compartments. We were surprised to discover that ORF6 exhibits a biased hydrophobic index and is predicted to be an amphipathic protein (**Figs. 2A** and **2B**). From these observations, we postulated two models to explain how ORF6 could be amphipathic and localize to membranous compartments. First, ORF6 is a transmembrane protein that forms higher order homo-or hetero-oligomers that shield the hydrophilic surface from the hydrophobic membrane environment. Second, ORF6 is a peripheral-membrane protein that orients the hydrophilic helical surfaces toward the cytoplasm and buries the hydrophobic portion within membrane surfaces.

**Figure 2.**
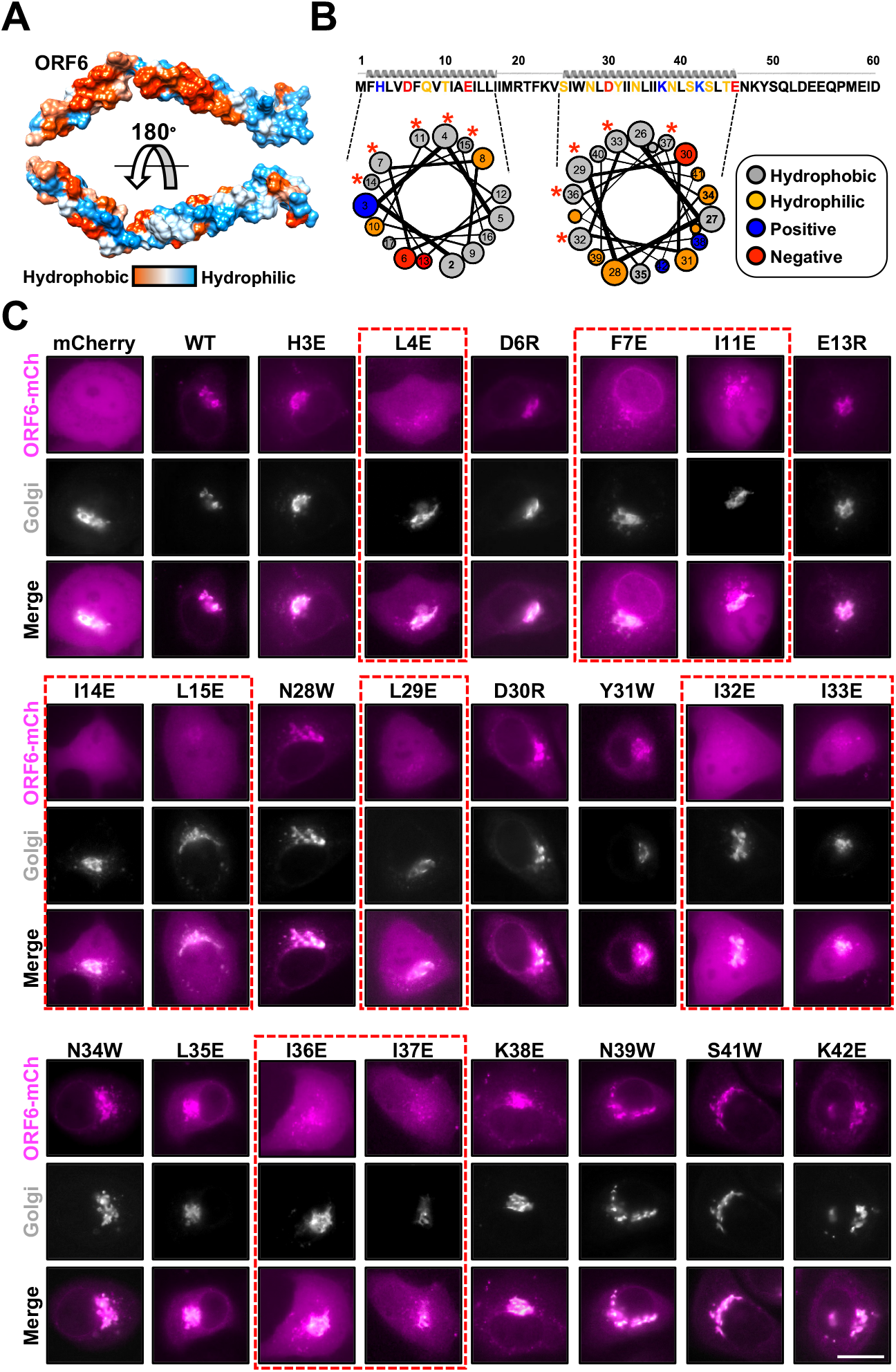
Putative amphipathic helices regulate ORF6 localization. (**A**) Surface representation of an ORF6 computational model depicting the hydrophobic surface index. (**B**) Helical wheel diagrams highlighting the amino acid composition of putative ORF6 helices. Positions are colored based on amino acid properties and asterisks indicate positions where mutations disrupted localization. (**C**) Representative live cell fluorescence microscopy images of the indicated mCherry-tagged proteins expressed in HeLa cells. Dashed red boxes highlight mutants that disrupt localization.

To explore these models, we generated and tested single amino acid substitution mutants that exchanged the wild-type residue for one with opposing biophysical properties on the respective helical surfaces. As depicted in **Figs. 2B** and **2C**, polar residues were replaced with tryptophan, charged residues were exchanged with opposing charge, hydrophobic residues were replaced with glutamate, and localization was assessed. If ORF6 is a transmembrane protein that requires intramembrane oligomerization, we postulated that substitutions made to either surface would disrupt localization. However, if ORF6 is a peripheral-membrane protein, only substitutions made to the hydrophobic membrane-interacting surface would exhibit disrupted localization. In support of the latter scenario, ORF6 mutants with substitutions on the hydrophilic surface exhibited normal localization; however, all ORF6 mutants with substitutions on the hydrophobic surface mis-localized (**Figs. 2B**, red asterisks; and **2C**, dashed boxes). Taken together, these results suggest that SARS-CoV-2 ORF6 is likely a peripheral-membrane protein that maintains steady-state localization through the ^18^IMRTFKV^24^ region and this contiguous hydrophobic surface.

To further explore the possibility that SARS-CoV-2 is a peripheral-membrane protein, we examined localization patterns in the presence of brefeldin A (BFA) and cycloheximide (CHX). Brefeldin A is a well characterized fungal toxin that promotes Golgi disassembly by inhibiting trafficking between the Golgi and ER compartments, subsequently resulting in the rapid redistribution of Golgi proteins to the ER (Lippincott-Schwartz et al., 1989). We reasoned if ORF6 peripherally associates with Golgi membranes, BFA treatment would induce cytoplasmic accumulation of ORF6, whereas if it’s a transmembrane protein, it would relocalize to the ER. The inclusion of the protein translation inhibitor CHX allows for tracking of steady-state ORF6 protein rather than observing newly synthesized protein that may artificially accumulate in the cytoplasm due to loss of the Golgi apparatus. As depicted in **Fig. 3A**, HeLa cells treated with BFA exhibited loss of Golgi stacks and subsequent redistribution of the Golgi marker to the ER. Importantly, BFA treatment induced SARS-CoV-2 ORF6 puncta accumulation in the cytoplasm, which resembled the 18-24Ala mutant that lacks the Golgi retention motif (**Figs. 1C** and **3A**). Interestingly, we did not observe the same phenomenon for SARS-CoV-1 ORF6 which fully redistributed to the ER, also mimicking its cognate 18-24Ala mutant (**Figs. 1C** and **3A**). Next, we wanted to determine if SARS-CoV-2 ORF6 puncta could re-associate with nascent Golgi membranes following BFA washout. As shown in **Fig. 3B**, HeLa cells imaged following BFA washout exhibited redistribution of the Golgi marker from the ER to discrete puncta in the cytoplasm, which were most likely reassembling Golgi stacks. Likewise, SARS-CoV-2 ORF6 colocalized with these Golgi puncta, whereas SARS-CoV-1 ORF6 remained colocalized with the ER following washout, suggesting that the cytoplasmic SARS-CoV-2 ORF6 proteins could more readily associate with nascent Golgi stacks (**Fig. 3B**). To further confirm BFA treatment induced redistribution of SARS-CoV-2 ORF6 proteins to the cytoplasm, we treated cells with BFA and then with digitonin. Digitonin is a non-ionic detergent that selectively permeabilizes the plasma membrane while leaving other membranous compartments intact. If SARS-CoV-2 ORF6 is indeed released from the Golgi membrane following BFA treatment, we hypothesized that ORF6 would deplete from the cytoplasm following digitonin treatment. First, we assessed the behavior of cells coexpressing mCherry and the Golgi marker to ensure digitonin treatment would only disrupt mCherry accumulation and not the Golgi apparatus. As expected, digitonin treatment, but not BFA treatment, resulted in a significant depletion of mCherry fluorescence in transfected cells as indicated by fluorescence microscopy and flow cytometry (**Fig. 3C**). Furthermore, the Golgi marker only exhibited differential distribution in the presence of BFA, not digitonin, indicating that the detergent was not grossly disrupting the structure of the Golgi. When HeLa cells expressing SARS-CoV-2 ORF6 were treated with digitonin alone, no impact on protein abundance or localization was observed (**Fig. 3C**). However, when cells were treated with BFA and then with digitonin, we observed significantly decreased fluorescence intensity for ORF6 but not the Golgi marker (**Fig. 3C**). Taken together, these observations combined with the mutational data in **Figs. 1** and **2** strongly support the model that SARS-CoV-2 ORF6 is a peripheral-membrane protein, and likely explains how some variants can non-specifically associate with membranous compartments (*i*.*e*., 18-24, RK-EE, and 24-47 mutants).

**Figure 3.**
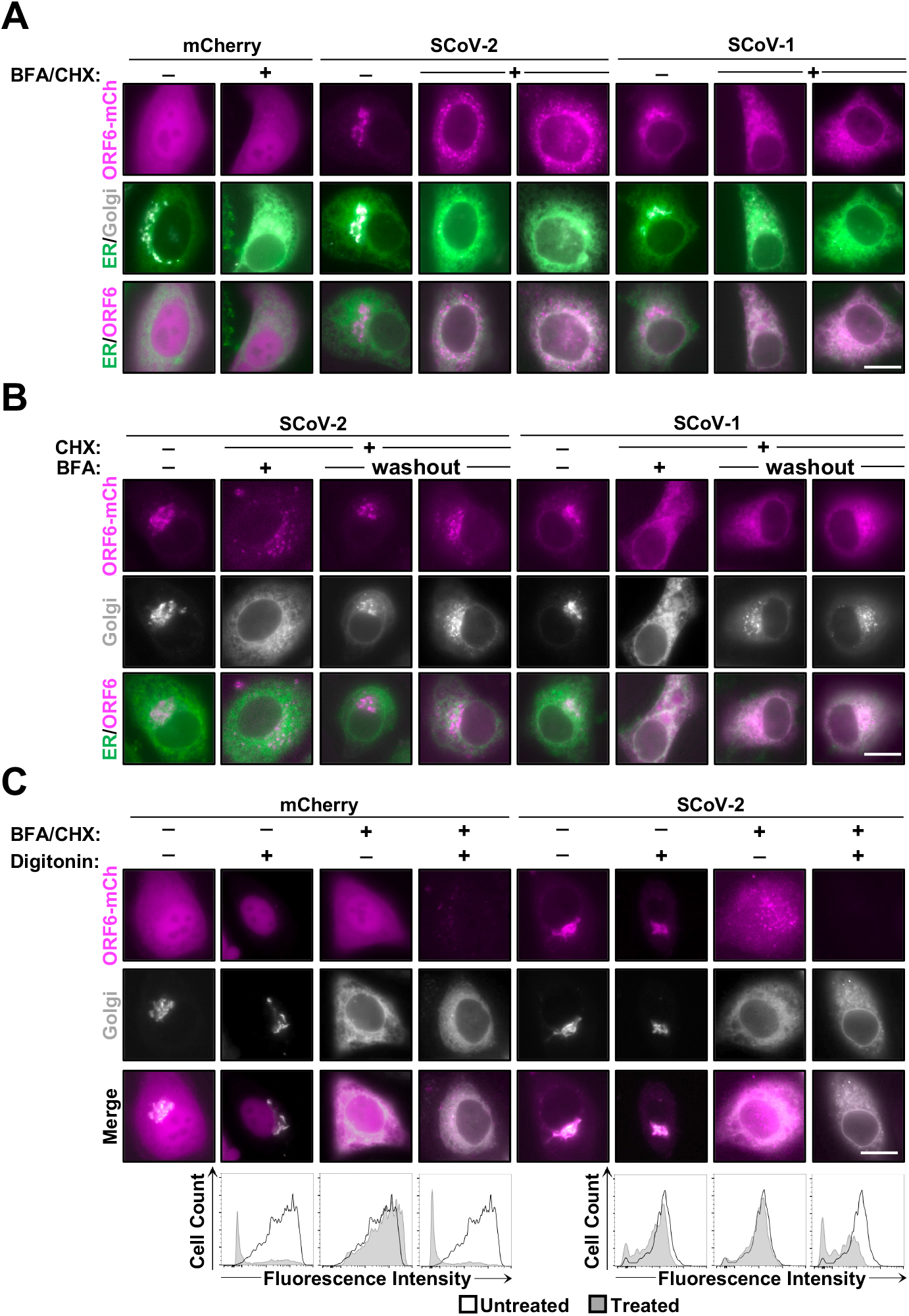
SARS-CoV-2 ORF6 is likely a peripheral membrane protein. (**A** and **B**) Representative live cell fluorescence microscopy images of the indicated mCherry-tagged proteins co-expressed in HeLa cells with ER and Golgi markers. Samples were either untreated or treated with indicated combinations of brefeldin A (BFA) and cycloheximide (CHX). (**C**) (Top) Representative live cell fluorescence microscopy images of either untreated or treated cells expressing the indicated mCherry-tagged proteins co-expressed in HeLa cells with a Golgi marker. (Bottom) Representative flow cytometry histograms displaying fluorescence intensity from HeLa cells expressing the indicated protein following treatment, or no treatment, with the indicated compounds.

### Membrane association of ORF6 is not required for interferon antagonism

ORF6 inhibits interferon synthesis and signaling by blocking nuclear translocation of activated STAT1 and IRF3 transcription factors through a direct interaction with Nup98-Rae1 (Frieman et al., 2007; Kato et al., 2021; Kopecky-Bromberg et al., 2007; Lei et al., 2020; Miorin et al., 2020; Xia et al., 2020). The most likely scenario is that ORF6 accumulation at the nuclear envelop facilitates a direct interaction with Nup98-Rae1 to block nuclear trafficking; however, this relationship has yet to be explored experimentally. To determine if membrane association is required for inhibiting nuclear trafficking, we examined a panel of localization disrupted mutants for their ability to block eGFP-KPNA2 trafficking. To ensure that ORF6 function was not altered due to the presence of the C-terminal mCherry tag, we generated a mCherry-T2A-ORF6 “self-cleaving” expression cassette, which allows for efficient detection of transfected cells without having to epitope-tag ORF6 (Salamango et al., 2019). For these experiments, we stably introduced eGFP-KPNA2 in the type II alveolar lung epithelial cell line A549 to test ORF6 activity in a more physiologically relevant cell model. Remarkably, all mutants tested were able to induce cytoplasmic accumulation of eGFP-KPNA2 at efficiencies comparable to wild-type (representative images in **Fig. 4A**, quantification in **Fig. 4C**). Next, we wanted to confirm these mutants also block nuclear accumulation of activated STAT1. To probe this activity, we treated A549 cells with type I IFN and assessed STAT1 localization in the presence or absence of ORF6 proteins. As anticipated, all mutants could inhibit nuclear accumulation of STAT1 at efficiencies comparable to wild-type (representative images in **Fig. 4B**, quantification in **Fig. 4D**). Furthermore, we assessed the ability of these mutants to inhibit interferon signaling using an IFNβ-eGFP reporter construct. We transiently co-expressed the IFNβ reporter with an activating RIG-I mutant (Mibayashi et al., 2007) in the presence or absence of the indicated ORF6 proteins and assessed eGFP expression. As depicted in **Fig. 4E**, both wild-type and mutant ORF6 proteins could efficiently suppress eGFP expression following stimulation with the activating RIG-I mutant. Lastly, to further confirm that membrane association is not required for inhibition of nuclear trafficking, we generated an inhibitor construct by fusing 3x copies of the ORF6/Nup98 interaction domain to mCherry and assessed eGFP-KPNA2 relocalization. As depicted in **Figs. 4F** and **4G**, expression of this construct was sufficient to drive cytoplasmic accumulation of eGFP-KPNA2.

**Figure 4.**
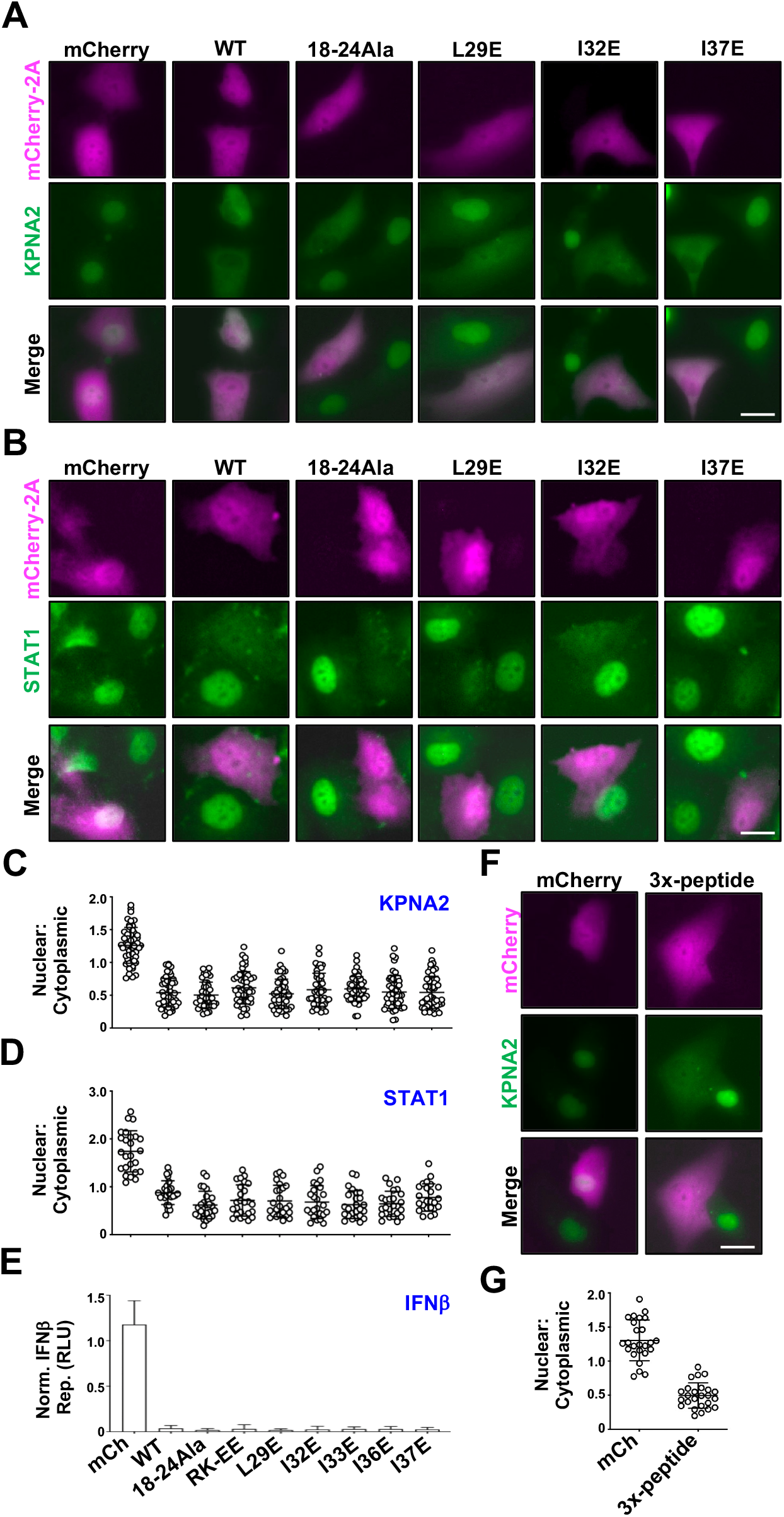
Membrane association of ORF6 is not required for interferon antagonism. (**A** and **B**) Representative live cell fluorescence microscopy images of the indicated mCherry-tagged proteins expressed in A549 cells stably expressing eGFP-KPNA2 (**A**) or following IFN-treatment and STAT1 immuno-labeling (**B**). (**C** and **D**) Quantification of localization patterns for the indicated constructs in A549 cells either stably expressing eGFP-KPNA2 or following IFN-treatment and STAT1 immuno-labeling. (**E**) Quantification of fluorescence from an IFNβ-eGFP reporter plasmid co-expressed with the indicated ORF6 proteins following RIG-I stimulation. (**F**) Representative live cell fluorescence microscopy images of A549 cells expressing either mCherry or the mCherry-3x-peptide construct. (**G**) Quantification of localization patterns for the indicated constructs in A549 cells.

Based on our findings here, it is evident that ORF6 localization is independent from IFN antagonism and raises the possibility that ORF6 may have additional functions within membrane networks to enhance viral replication.

## MATERIALS AND METHODS

### Cell culture and cloning

A549, HEK293FT, and HeLa cells were maintained in RPMI (Hyclone, South Logan, UT) supplemented with 10% FBS (Gibco, Gaithersburg, MD) and 0.5% pen/strep (50 units). Cells were transfected with PEI using a ratio of 3 µL per 1 µg of DNA. To generate the A549 eGFP-KPNA2 stable cell line, roughly 250,000 HEK293FT cells in a 6-well plate were transfected with a pQCXIH retroviral vector containing the eGFP-KPNA2 expression cassette, an MLV-GagPol packaging vector, and a VSV-G vector. Media was harvested 48 hours post-transfection and frozen at −80°C for 4-6 hours, thawed and centrifuged at 1500 x g, and overlaid on A549 cells. To generate a pure cell population, cells were treated with hygromycin B (Sigma, 500 µg/ml) 48 hours post-transduction. All ORF6 mutants were generated by PCR amplification using Phusion high fidelity DNA polymerase (NEB, Ipswich, MA) and overlapping PCR to introduce the indicated mutations. To generate the peptide inhibitor construct, we fused 3x tandem copies of the ORF6/Nup98 interaction domain separated by 12 amino acid glycine/serine/threonine linkers (repeat peptide sequence: KYSQLDEEQPMEID) to the C-terminus of mCherry. The IFNβ reporter construct was generated by cloning an IFN responsive promoter element upstream of eGFP in the pcDNA-5TO expression vector and the constitutively active RIG-I mutant vector was generated by cloning RIG-I amino acids 1-242 into an mTagBFP2-T2A-expression cassette in a lentiviral vector. All constructs were confirmed by restriction digestion and Sanger sequencing.

For experiments using the IFNβ and RIG-I expression constructs, roughly 250,000 A549 cells were seeded into 12 well culture plates and allowed to adhere overnight. The next day, cells were co-transfected with the indicated combinations of IFNβ-eGFP reporter, activating RIG-I, and ORF6 expression plasmids at 100 ng DNA per construct. At 48 hours post-transfection, eGFP fluorescence was quantified (ImageJ software) and then graphed with Prism 6.0 (GraphPad Software).

### Fluorescence microscopy and immunostaining

All localization and immunostaining experiments were repeated at least 3 independent times and representative images are depicted from surveying at least 5 fields of view from each condition with at least 25-30 cells exhibiting similar phenotypes (all scale bars shown are at 10 µm). In addition, localization experiments were carried out using HeLa cells as they are routinely used for microscopy experiments and localization studies due to their morphological characteristics. Roughly 6,500 HeLa cells were seeded into an 8-well #1.5 glass bottom chamber slide (Ibidi #80826) and transfected with 100 ng of the indicated ORF6 expression construct along with either 50 ng of an ER marker (eGFP-Calnexin; Addgene: 57122), 50 ng of a Golgi marker [mTag-β-galactosidase (1-61)], or 50 ng of a mitochondrial marker (eBFP2-Mito, Addgene: 55248). The next day, cells were imaged using a 60x oil immersion objective on an EVOS M5000 fluorescence microscope.

Prior to immuno-labeling and imaging of STAT1, A549 cells were treated with 3000 U/mL type I interferon for 45 minutes. Immuno-labeling of STAT1 was performed as follows: 24 hours post transfection, cells were washed with PBS and fixed in 4% PFA at room temperature for 10 minutes. Following fixation, cells were washed using PBS+0.3% Triton X-100 (PBST) three times in 5-minute intervals and then blocked using PBST supplemented with 5% BSA, 10% goat serum, and 0.3 M glycine for 1 hour at room temperature. After blocking, samples were incubated with primary anti-STAT1 antibody (Cell Signaling Technology, 9172) in PBST with 5% BSA overnight at 4 degrees. The next day, samples were washed 3 times with PBS and then incubated with secondary anti-Rabbit-AlexaFluor488 (Cell Signaling Technology, 4412) and anti-mCherry-AlexaFluor594 (Invitrogen #M11240) antibodies in PBS with 5% BSA for 1 hour at room temperature. Finally, cells were washed 3 times with PBS and then imaged at 60x on an EVOS M5000 fluorescence microscope. For quantification of nuclear fluorescence, individual cells expressing the indicated ORF6 proteins were scored for KPNA2 or STAT1 localization by dividing the nuclear fluorescence intensity by the cytoplasmic fluorescence intensity (n=50 for KPNA2 and n=25 for STAT1), and then graphed with Prism 6.0 (GraphPad Software). Subcellular compartments were defined based on DAPI staining (nucleus).

### Chemical inhibitor treatments

For BFA/CHX inhibitor treatments, culture media was replaced with complete media supplemented with 5 µM BFA and 2 µM CHX and allowed to incubate for 30 minutes prior to imaging. For inhibitor washout, samples were rinsed with PBS following an initial 30-minute BFA/CHX incubation and then complete media supplemented with 2 µM CHX was added. Cells were allowed to recover for 30 minutes before imaging reconstitution of the Golgi apparatus. For digitonin treatments, transfected cells were either left untreated, treated with 50 µg digitonin for 10 minutes, treated with 5 µM BFA/2 µM CHX for 30 minutes, or, treated with 5 µM BFA/2 µM CHX for 30 minutes and then with 50 µg digitonin for 10 minutes.

### Flow cytometry

All flow cytometry experiments were repeated 3 independent times and representative histograms are depicted from one experiment. Quantification of fluorescence intensity was performed using a Becton Dickinson FACScan flow cytometer. Briefly, roughly 125,000 HeLa cells were seeded into a 12 well plate and transfected 24 hours after plating with 300 ng of the indicated ORF6/fluorescent protein DNA and 200 ng of DNA for the Golgi marker. The next day, cells were treated as described above with the following exception: digitonin treatment was not performed until after cells were removed from culture plates to better preserve cell integrity. Following inhibitor treatment, cells were removed from plates using 0.025% Trypsin/EDTA solution, centrifuged at 300 x g for 5 minutes, and then re-suspended in 2% FBS + 5 µM BFA/2µM CHX in PBS. At this point, 50 µg of digitonin was added for 10 minutes and samples were subjected to flow cytometry and analyzed using FloJo software.

## ACKNOWLEDGEMENTS

We thank Drs. Erich Mackow, Nancy Reich-Marshall, and Patrick Hearing for intellectual discussions and constructive feedback. This work was supported by startup funds provided by the Department of Microbiology and Immunology and Renaissance School of Medicine at Stony Brook University. All authors declare no conflict of interest.

## Notes

### Competing Interest Statement

The authors have declared no competing interest.

## REFERENCES

Addetia, A., Lieberman, N. A. P., Phung, Q., Hsiang, T. Y., Xie, H., Roychoudhury, P., Shrestha, L., Loprieno, M. A., Huang, M. L., Gale, M. et al. (2021). SARS-CoV-2 ORF6 Disrupts Bidirectional Nucleocytoplasmic Transport through Interactions with Rae1 and Nup98. mBio 12, e00065–21.

Banfield, D. K. (2011). Mechanisms of protein retention in the Golgi. Cold Spring Harb Perspect Biol 3, a005264.

Frieman, M., Yount, B., Heise, M., Kopecky-Bromberg, S. A., Palese, P. and Baric, R. S. (2007). Severe acute respiratory syndrome coronavirus ORF6 antagonizes STAT1 function by sequestering nuclear import factors on the rough endoplasmic reticulum/Golgi membrane. J Virol 81, 9812–24.

Gordon, D. E. Jang, G. M. Bouhaddou, M. Xu, J. Obernier, K. White, K. M. O’Meara, M. J. Rezelj, V. V. Guo, J. Z. Swaney, D. L. et al. (2020). A SARS-CoV-2 protein interaction map reveals targets for drug repurposing. Nature 583, 459–468.

Gunalan, V., Mirazimi, A. and Tan, Y. J. (2011). A putative diacidic motif in the SARS-CoV ORF6 protein influences its subcellular localization and suppression of expression of co-transfected expression constructs. BMC Res Notes 4, 446.

Hu, L., Li, L., Xie, H., Gu, Y. and Peng, T. (2011). The Golgi localization of GOLPH2 (GP73/GOLM1) is determined by the transmembrane and cytoplamic sequences. PLoS One 6, e28207.

Huang, C., Peters, C. J. and Makino, S. (2007). Severe acute respiratory syndrome coronavirus accessory protein 6 is a virion-associated protein and is released from 6 protein-expressing cells. J Virol 81, 5423–6.

Kato, K., Ikliptikawati, D. K., Kobayashi, A., Kondo, H., Lim, K., Hazawa, M. and Wong, R. W. (2021). Overexpression of SARS-CoV-2 protein ORF6 dislocates RAE1 and NUP98 from the nuclear pore complex. Biochem Biophys Res Commun 536, 59–66.

Kopecky-Bromberg, S. A., Martínez-Sobrido, L., Frieman, M., Baric, R. A. and Palese, P. (2007). Severe acute respiratory syndrome coronavirus open reading frame (ORF) 3b, ORF 6, and nucleocapsid proteins function as interferon antagonists. J Virol 81, 548–57.

Kumar, P., Gunalan, V., Liu, B., Chow, V. T., Druce, J., Birch, C., Catton, M., Fielding, B. C., Tan, Y. J. and Lal, S. K. (2007). The nonstructural protein 8 (nsp8) of the SARS coronavirus interacts with its ORF6 accessory protein. Virology 366, 293–303.

Lee, J. G., Huang, W., Lee, H., van de Leemput, J., Kane, M. A. and Han, Z. (2021). Characterization of SARS-CoV-2 proteins reveals Orf6 pathogenicity, subcellular localization, host interactions and attenuation by Selinexor. Cell Biosci 11, 58.

Lei, X., Dong, X., Ma, R., Wang, W., Xiao, X., Tian, Z., Wang, C., Wang, Y., Li, L., Ren, L. et al. (2020). Activation and evasion of type I interferon responses by SARS-CoV-2. Nat Commun 11, 3810.

Lippincott-Schwartz, J., Yuan, L. C., Bonifacino, J. S. and Klausner, R. D. (1989). Rapid redistribution of Golgi proteins into the ER in cells treated with brefeldin A: evidence for membrane cycling from Golgi to ER. Cell 56, 801–13.

Mibayashi, M., Martínez-Sobrido, L., Loo, Y. M., Cárdenas, W. B., Gale, M. and García-Sastre, A. (2007). Inhibition of retinoic acid-inducible gene I-mediated induction of beta interferon by the NS1 protein of influenza A virus. J Virol 81, 514–24.

Miorin, L., Kehrer, T., Sanchez-Aparicio, M. T., Zhang, K., Cohen, P., Patel, R. S., Cupic, A., Makio, T., Mei, M., Moreno, E. et al. (2020). SARS-CoV-2 Orf6 hijacks Nup98 to block STAT nuclear import and antagonize interferon signaling. Proc Natl Acad Sci U S A 117, 28344–28354.

Netland, J., Ferraro, D., Pewe, L., Olivares, H., Gallagher, T. and Perlman, S. (2007). Enhancement of murine coronavirus replication by severe acute respiratory syndrome coronavirus protein 6 requires the N-terminal hydrophobic region but not C-terminal sorting motifs. J Virol 81, 11520–5.

O’Keefe, S., Roboti, P., Duah, K. B., Zong, G., Schneider, H., Shi, W. Q. and High, S. (2021). Ipomoeassin-F inhibits the in vitro biogenesis of the SARS-CoV-2 spike protein and its host cell membrane receptor. J Cell Sci 134.

Salamango, D. J., Ikeda, T., Moghadasi, S. A., Wang, J., McCann, J. L., Serebrenik, A. A., Ebrahimi, D., Jarvis, M. C., Brown, W. L. and Harris, R. S. (2019). HIV-1 Vif Triggers Cell Cycle Arrest by Degrading Cellular PPP2R5 Phospho-regulators. Cell Rep 29, 1057–1065.e4.

Silvas, J. A., Vasquez, D. M., Park, J. G., Chiem, K., Allué-Guardia, A., Garcia-Vilanova, A., Platt, R. N., Miorin, L., Kehrer, T., Cupic, A. et al. (2021). Contribution of SARS-CoV-2 Accessory Proteins to Viral Pathogenicity in K18 Human ACE2 Transgenic Mice. J Virol 95, e0040221.

Tu, L., Chen, L. and Banfield, D. K. (2012). A conserved N-terminal arginine-motif in GOLPH3-family proteins mediates binding to coatomer. Traffic 13, 1496–507.

Wang, J., Chen, J., Enns, C. A. and Mayinger, P. (2013). The first transmembrane domain of lipid phosphatase SAC1 promotes Golgi localization. PLoS One 8, e71112.

Wang, P., Ye, Z. and Banfield, D. K. (2020). A novel mechanism for the retention of Golgi membrane proteins mediated by the Bre5p/Ubp3p deubiquitinase complex. Mol Biol Cell 31, 2139–2155.

Xia, H., Cao, Z., Xie, X., Zhang, X., Chen, J. Y., Wang, H., Menachery, V. D., Rajsbaum, R. and Shi, P. Y. (2020). Evasion of Type I Interferon by SARS-CoV-2. Cell Rep 33, 108234.

Zhao, J., Falcón, A., Zhou, H., Netland, J., Enjuanes, L., Pérez Breña, P. and Perlman, S. (2009). Severe acute respiratory syndrome coronavirus protein 6 is required for optimal replication. J Virol 83, 2368–73.

Zhou, H., Ferraro, D., Zhao, J., Hussain, S., Shao, J., Trujillo, J., Netland, J., Gallagher, T. and Perlman, S. (2010). The N-terminal region of severe acute respiratory syndrome coronavirus protein 6 induces membrane rearrangement and enhances virus replication. J Virol 84, 3542–51.

